# CreER activation transiently impairs angiogenesis by slowing endothelial proliferation

**DOI:** 10.1101/2025.05.22.655564

**Authors:** Elena Ioannou, Mengmeng Dong, Victoria S. Rashbrook, Christiana Ruhrberg, Martina Rudnicki

## Abstract

Tamoxifen-inducible gene targeting in mice with estrogen receptor-dependent Cre (CreER) recombinase has enormously advanced vascular biology research. However, CreER activation under the control of vascular endothelial promoters is now recognized to cause off-target effects that impair angiogenesis in the widely used perinatal mouse retina model. Although ubiquitously expressed CreER is also used to study retinal angiogenesis, it remains unknown whether it causes similar or more severe off-target effects compared to endothelial-selective CreER activation. Moreover, the cellular processes disrupted by CreER-induced endothelial toxicity remain to be identified. Here, we demonstrate that ubiquitous CreER activation in postnatal mice decreases body growth throughout the period of retinal angiogenesis and impairs retinal angiogenesis in a tamoxifen dose-dependent manner. We further show that CreER activation from both endothelial and ubiquitously expressed CreER transgenes suppresses endothelial cell proliferation downstream of p21/CDKNA1 upregulation. By contrast, we find that p21/CDKNA1 is not upregulated in quiescent adult retinal endothelium, and that CreER-induced postnatal angiogenesis defects recover two weeks after tamoxifen discontinuation. Altogether, our findings indicate that ubiquitous promoters should be avoided for CreER expression when studying endothelial genes, and that short-term retinal angiogenesis studies require endothelial CreER toxicity controls that may be less critical for adult vascular studies.

## Introduction

Cre recombinase-mediated gene ablation offers an invaluable genetic tool to uncover molecules that regulate vascular biology in the mouse, the major mammalian model system [1]. Commonly, this method involves the use of Cre fused to a human estrogen receptor variant (ER); this fusion protein, termed CreER, is retained in the cytoplasm until it binds 4-hydroxy-tamoxifen, thereby enabling its nuclear translocation to induce gene recombination [2]. Administering 4-hydroxy-tamoxifen or its precursor tamoxifen to promote Cre-mediated recombination therefore allows to control the time of deletion for genes that are flanked by loxP sites (floxed). By activating CreER after birth in vascular endothelial cells, this method circumvents the lethality caused by ablating essential vascular genes *in utero* [3]. Accordingly, this method allows to uncover molecular pathways that contribute to the growth, remodeling and function of postnatal blood vessels [1,4]. For example, the CreER system has been used to demonstrate the role of VEGF and notch signalling in postnatal angiogenesis [5–11].

Despite its usefulness for genetic investigation, a prior study demonstrated that CreER activation in endothelial cells with the commonly used *Cdh5*-CreER [11] or *Pdgfb*-CreER [12] transgenes can impair angiogenesis [13]. However, it has not yet been addressed whether ubiquitously expressed transgenes such as *Cagg*-CreER or *Rosa26*-CreER cause similar or worse CreER toxicity during angiogenesis, even though these transgenes have also been used to delete genes in endothelial cells or cells in their environment (e.g., [14–16,9]). Moreover, the mechanism that underlies CreER toxicity to impair angiogenesis remain unknown. Prior studies in non-endothelial cell types have suggested that Cre recombinase has toxic effects because it recognizes endogenous genomic DNA sequences that resemble the engineered loxP sequences used to mark genes for recombination [17–19]. Thus, off-target endonuclease activity directed at pseudo-loxP sites has been linked to DNA breaks, chromosomal aberrations, reduced cell proliferation, increased apoptosis and pro-inflammatory responses in non-endothelial cell types [20,21,4]. However, whether CreER activation also perturbs endothelial cell health in this manner has not been addressed. Moreover, it remains unknown whether the toxic effects of endothelial CreER activation are restricted to angiogenic endothelium or similarly affect the quiescent adult endothelium.

Considering that the mouse perinatal retina is widely used to identify molecules that regulate postnatal angiogenesis [22–24], we have used this model to investigate the emergence, mechanisms and resolution of CreER toxicity in endothelial cells. In the mouse, blood vessels emerge from vessels in the optic nerve head shortly after birth to form a superficial vascular plexus in the retina. Led by filopodia studded tip cells, vessels grow forwards to cover the retina and fuse laterally, thereby establishing a planar network that is exquisitely suited for high-resolution imaging and quantification of angiogenic processes [24,22,25,3]. Prior work showed that CreER activation in endothelial cells impaired both vessel extension and branching to significantly reduce vascular complexity as key angiogenic parameters in the retina [13]. Here, we show that CreER activation (but not tamoxifen or the vehicle used to administer tamoxifen) impairs retinal angiogenesis downstream of cell cycle stalling. Upregulation of the p21 cyclin dependent kinase inhibitor A1 (CDKNA1), which mediates both the TP53-induced and TP53-independent response to DNA damage, was the first indicator of CreER toxicity in endothelial cells. By contrast, CreER activation did not cause p21 upregulation or reduce the density of quiescent adult retinal vasculature. Our findings inform on the confounding mechanisms that might skew data interpretation when using the CreER-loxP system to assess candidate angiogenesis regulators whilst also demonstrating that targeting of the adult vasculature does not induce similar off-target effects.

## Results

### Tamoxifen or its vehicle do not reduce weight gain or retinal angiogenesis in neonatal mice

Vegetable oils are commonly used as vehicle for tamoxifen administration but can cause inflammation in adult mice when delivered via intraperitoneal injection [26]. To determine whether the intraperitoneal injection of vegetable oil as a vehicle affects growth or angiogenesis in postnatal mice, we delivered two doses, one on perinatal day (P) 2 and one on P4, and examined both male and female C57Bl6/J wild type pups on P7 (**Fig. 1A**). We found that vegetable oil on its own did not affect pup weight gain or retinal vascular network extension when compared to untreated pups (**Fig. 1A,B**). As tamoxifen has known vascular effects [27] and can reduce body weight independently of CreER activation in adult mice [28], we also investigated the effect of a range of tamoxifen doses commonly used to activate CreER for retinal gene targeting in neonatal pups. For this experiment, we tested three tamoxifen doses commonly used for gene targeting in the neonatal period, 50, 100 or 150 µg (e.g., [14–16,29,30]). However, intraperitoneal injection of tamoxifen in vegetable oil on P2 and P4 did not decrease body weight or vascular extension in male or female C57Bl6/J pups on P7 (average body weight: 50 μg, 3.93 g; 100 μg, 3.94 g; 150 μg, 3.80 g; average vascular extension relative to retinal radius: 50 μg, 0.80; 100 μg, 0.87; 150 μg, 0.86).

**Fig. 1.**
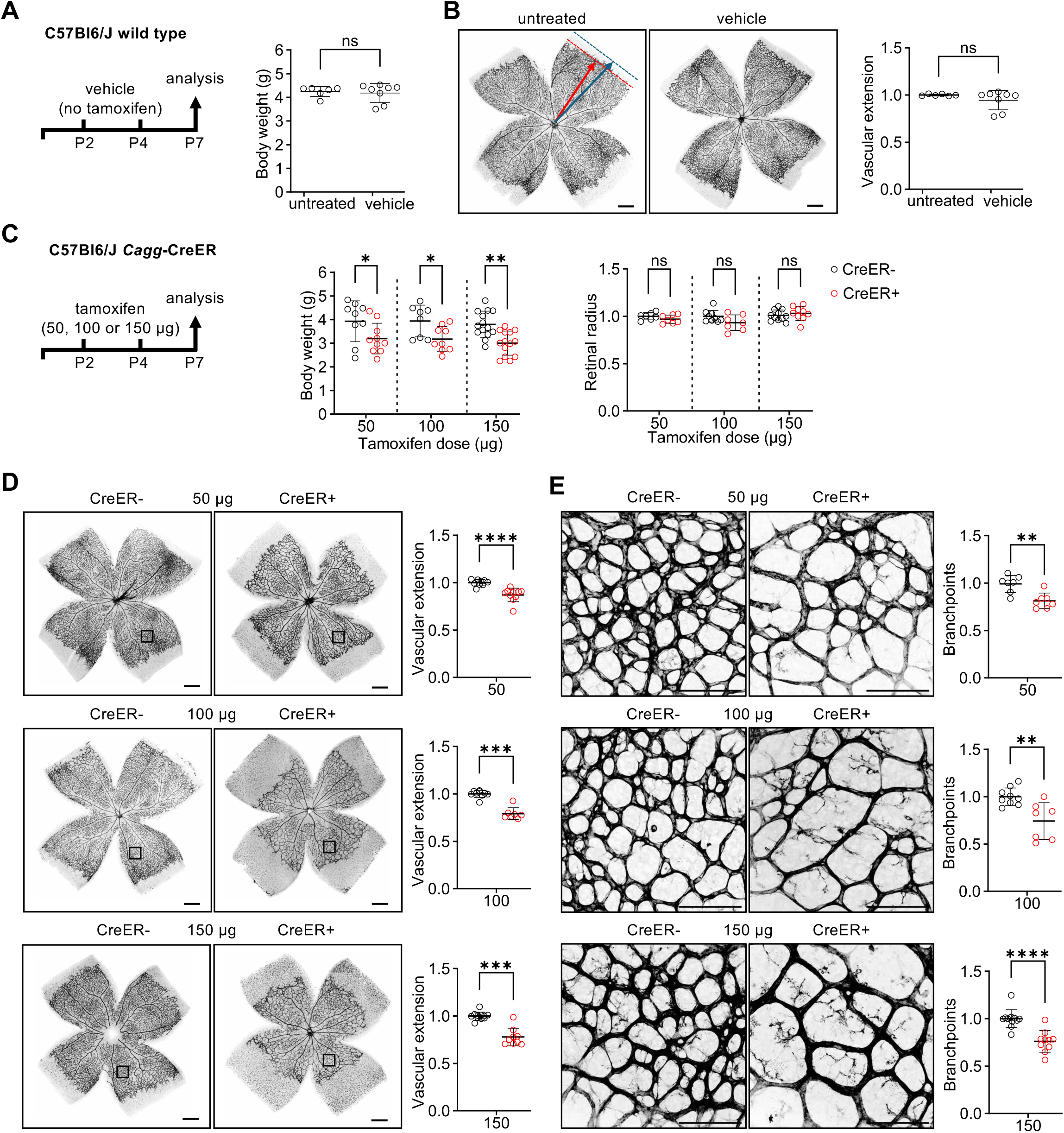
*Cagg*-CreER activation impairs weight gain and retinal angiogenesis in postnatal mice. (**A**) Schematic representation of the experimental setup and body weight measurement of P7 wild-type C57Bl6/J pups treated with vehicle (vegetable oil, n = 8) on postnatal day (P) 2 and P4, compared to untreated controls (n = 6). (**B**) Representative images of P7 whole-mount retinas stained with IB4 from untreated and vehicle-treated pups, and quantification of vascular extension (n = 6 retinas per group). Vascular extension was measured as the ratio of the distance from the optic nerve head to the peripheral vascular front (red arrow and line), relative to total retinal radius (blue arrow and dotted line). (**C-E**) Schematic representation of the tamoxifen treatment of *Cagg*-CreER+ and CreER-pups (**C**), along with body weight and retinal radius on P7 following tamoxifen administration at the indicated doses on P2 and P4 (n = 7-14 pups per group). (**D-E**) Representative images of P7 CreER+ and CreER-whole-mount retinas stained with IB4, including quantification of vascular extension (**D**, n = 7-11 pups per group) and branchpoints (**E**, n = 7-12 pups per group). Boxes in (**D**) indicate regions between an artery and a vein shown at higher magnification in (**E**) and used for branchpoint analysis. Scale bars: 500 µm (vascular extension); 100 µm (branchpoints). Data are shown as mean ± SD. Each data point represents one mouse (body weight) or the average of four measurements from one retina from one mouse (retinal radius and vascular parameters). Retinal data are normalized to untreated controls (**B**), or CreER-controls (**C-E**). \**P*< 0.05; \*\**P*< 0.01; \*\*\**P*<0.001; \*\*\*\**P*<0.001; ns: not significant (*P*>0.05). Mann-Whitney U test.

### Ubiquitous CreER activation reduces weight gain and retinal angiogenesis in neonatal mice

Next, we compared tamoxifen-injected neonatal C57BL6/J mice carrying or lacking the *Cagg-CreER* transgene [31] without floxed genes that could regulate angiogenesis to distinguish toxic effects of CreER activation from the effects of gene targeting. After delivering two doses of 50, 100 or 150 µg tamoxifen on P2 and P4 each, we examined pups on P7 (**Fig. 1C**). We found that male and female P7 pups with activated CreER (CreER+) had lower body weight compared to control mice lacking CreER (CreER-) across all tamoxifen doses examined (**Fig. 1C**). By contrast, the retina radius was similar in pups with or without activated CreER at all three doses examined (**Fig. 1C**), which allowed us to compare the effect of tamoxifen-induced CreER activation on retinal angiogenesis in the absence of confounding effects of eye size. Both vascular network extension and retinal vascular branch density, key angiogenic parameters, were significantly reduced in retinas of CreER+ compared to CreER-mice at all three tamoxifen doses examined (**Fig. 1D,E**). Although the reduction in vascular branchpoints was consistent across all three tamoxifen doses (*P* > 0.48), vascular extension was significantly lower with 50 μg tamoxifen compared to both 100 μg (*P* = 0.02) and 150 μg (*P* < 0.001; no significant difference observed between the 100 and 150 μg doses). Further, we found no evidence for treatment-by-sex interaction (χ² = 2.61, df = 2, *P* = 0.27), suggesting similar effects of CreER toxicity on retinal vascularisation in both sexes. Notably, the presence of loxP sites (introduced via a floxed *Rosa26^tdTom^*reporter cassette) did not ameliorate the detrimental effect of CreER activation on vascular extension (**Fig. S1A**).

Together, these findings demonstrate that a ubiquitously expressed CreER transgene used to target retinal angiogenesis, impairs retinal angiogenesis at the lowest dose examined and even when the genome contained engineered loxP sites as intended CreER targets. Additionally, ubiquitous *Cagg-CreER* activation impaired body growth, which was not seen with endothelial-selective CreER activation.

### CreER activation-induced retinal vascular defects recover over time

In adult mice, tamoxifen is metabolized to 4-OHT, whereby both compounds have a serum half-life of less than 10 hours and are barely detectable by 24 - 48 hours after administration [32,33]. We therefore examined whether tamoxifen discontinuation allowed body weight and retinal angiogenesis to recover from the defects caused by earlier tamoxifen-induced CreER activation after pups had received two doses of 100 μg tamoxifen on P2 and P4. This dose was chosen, because it disrupts both vascular extension and branching and has been used to examine gene deletion in multiple prior retinal angiogenesis studies. Then, we examined retinal vasculature at P21 two weeks after tamoxifen had been discontinued (**Fig. 2A**). At this time, tamoxifen-injected CreER+ mice still had significantly lower body weight compared to control littermates (**Fig. 2B**). We next acquired confocal z-projections that spanned the depth of all three vascular plexi of the maturing retinal vasculature (**Fig. 2C**) and grouped individual slices of the z-projection according to their representation of each plexus (**Fig. 2D**). Although the complexity of the superficial vascular plexus was impaired at P7 (see **Fig. 1E**), we observed normal vascular coverage and branchpoint complexity in the superficial retinal vascular plexus at P21 (**Fig. 2D**). Moreover, the deep plexus, which sprouts from the superficial plexus after P7, and the intermediate plexus, which forms from the deep plexus after P12 [23], also had normal vascular complexity (**Fig. 2D**). Accordingly, tamoxifen discontinuation allowed for recovery of a vascular plexus that was initially impaired by CreER toxicity. Additionally, we found that past CreER activation did not affect the growth of new vasculature after tamoxifen had been cleared. Together, these findings suggest that CreER activation induces vascular growth defects that can be compensated over time, whereas systemic consequences, such as reduced body weight, persist beyond the treatment period.

**Fig. 2.**
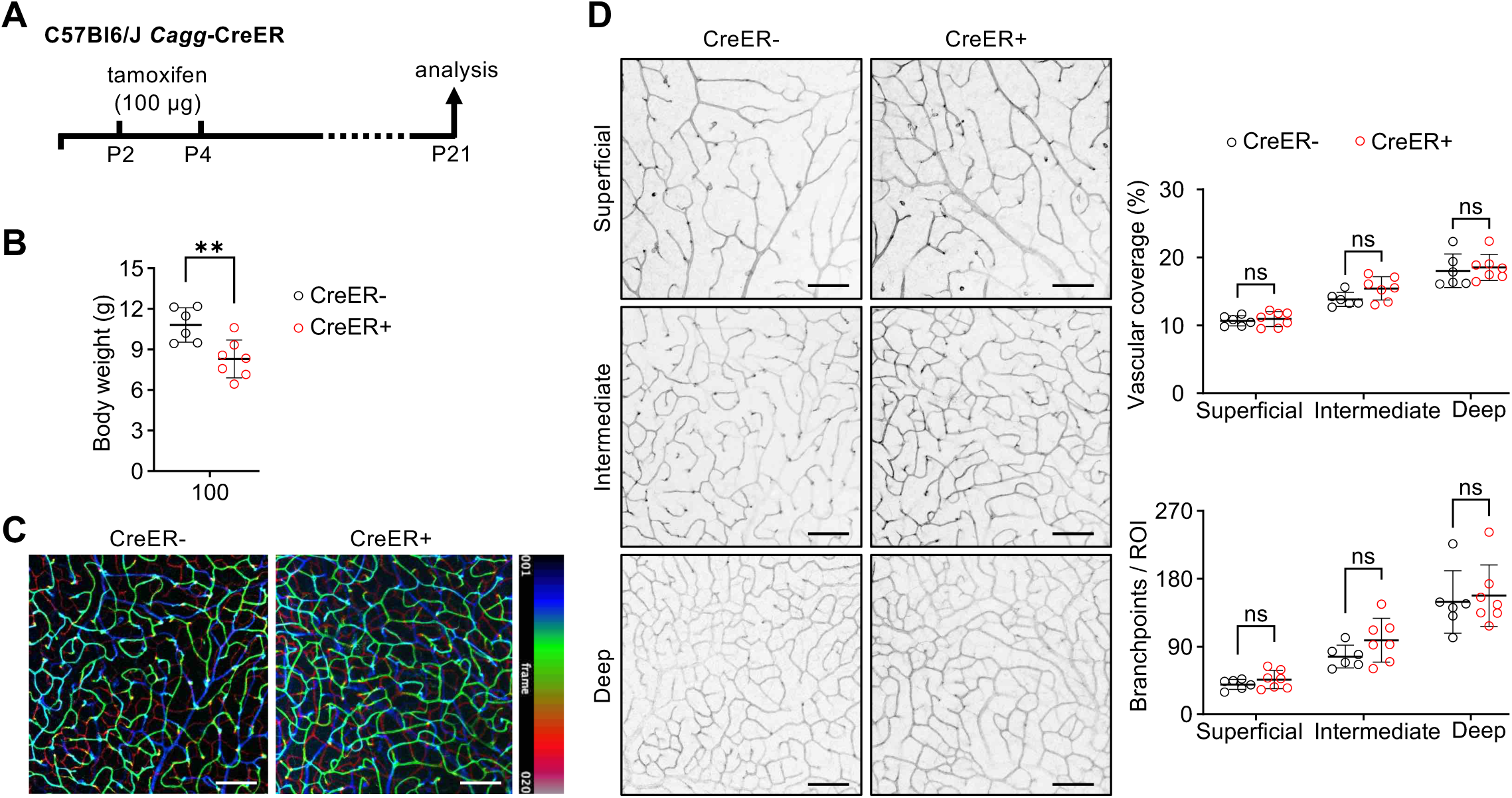
Recovery of retinal angiogenesis but not body weight defects caused by past CreER activation. (**A-D**) Schematic representation of the experimental setup (**A**) and body weight (**B**) of *Cagg*-CreER+ and CreER-littermate pups on a C57BL6/J background at P21 after treatment with 100 µg tamoxifen on P2 and P4 (n = 6-7 per group). **(C)** Representative projection of confocal z-stack images from whole-mount retinas of CreER+ and CreER-pups stained with IB4, with the vascular plexi resolved by pseudo-coloring according to retinal depth (20 z-slices; blue: superficial, green: intermediate, red: deep); **(D)** Representative images of the three IB4+ retinal plexi, shown in greyscale, extracted from the confocal z-projection shown in (**C**), and quantification of vascular coverage and branchpoints per region of interest (ROI) between an artery and vein for each plexus (n = 6-7 per group). Scale bars: 100 µm. Data are shown as mean ± SD. Each data point represents one mouse (body weight) or the average of four regions of interest (ROI) from one retina from one mouse (vascular measurements). \*\**P* < 0.01; ns: not significant (*P* > 0.05). Mann-Whitney U test.

### CreER activation impairs cell cycle progression during retinal angiogenesis

To understand the mechanisms underlying CreER-induced angiogenesis defects, we administered one or two doses of 100 μg tamoxifen and examined retinas 24 hours after the single or the second tamoxifen injection for altered endothelial cell behavior that might explain subsequent angiogenesis defects (**Fig. 3A**). Although the body weight of CreER- and CreER+ pups was still similar at this time (**Fig. S1B,C**), quantitative analysis of retinas stained for IB4 together with an antibody for the endothelial-specific transcription factor ERG [34] revealed 35% fewer ERG⁺ endothelial cells in CreER+ compared to CreER-retinas after the second tamoxifen dose, but not after a single tamoxifen dose or after vehicle treatment (**Fig. 3A**). Consistent with these observations, quantitative RT-PCR analysis of retinas from the opposing eye of the same mice showed a 30% decrease of *Erg* transcript levels after two but not one tamoxifen dose (**Fig. 3A**). These findings suggest that prolonged CreER activation reduces vascular growth.

**Fig. 3.**
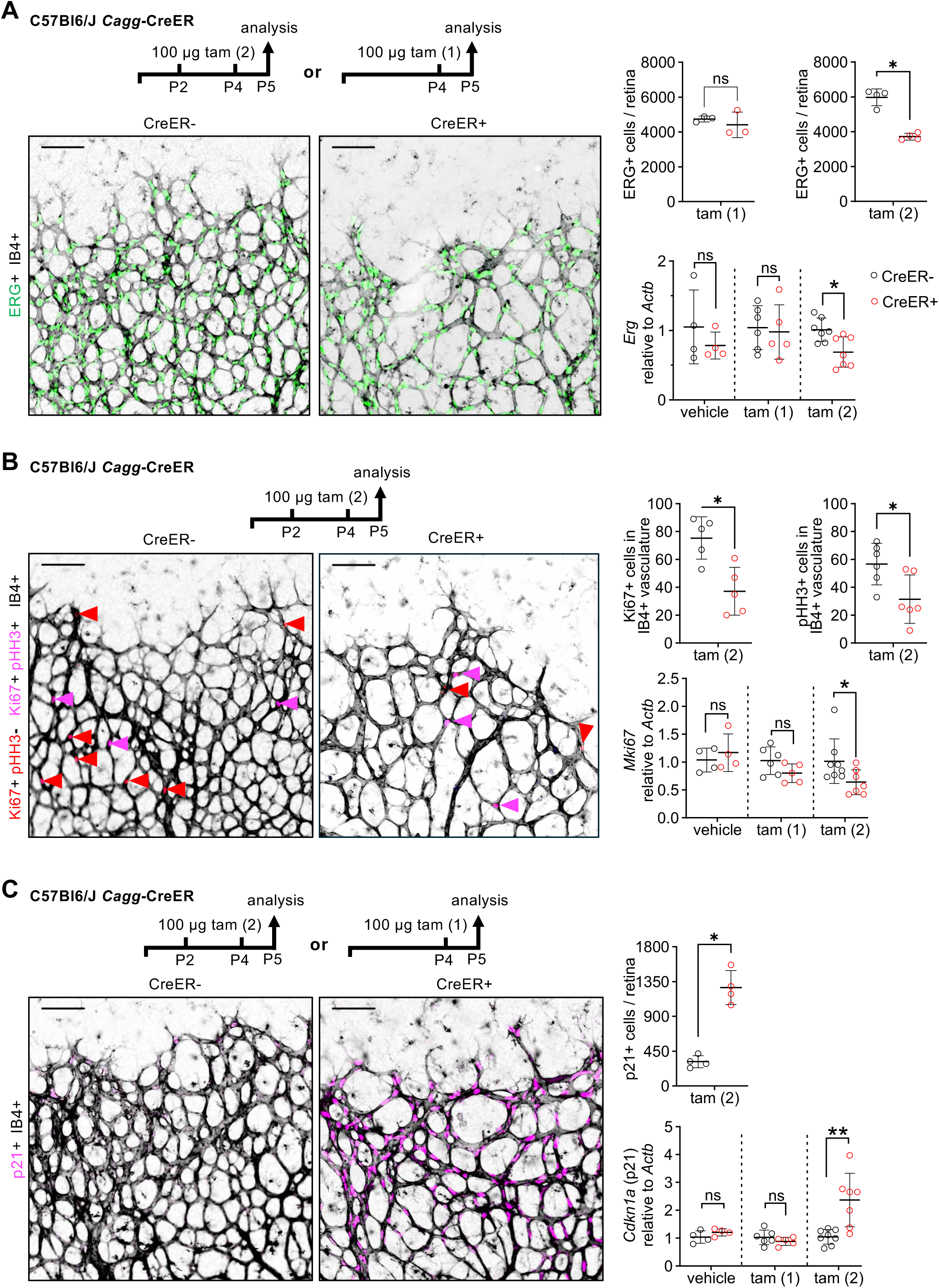
*Cagg*-CreER activation reduces endothelial cell proliferation and upregulates p21. (**A-C**) *Cagg*-CreER and CreER-littermate pups on a C57BL6/J background were treated with either vehicle, 100 µg tamoxifen on P2 and P4, referred to as the tam (2) condition, or 2 doses of vehicle, or with 100 µg tamoxifen on P4 only, referred to as the tam (1) condition, and then analyzed on P5; schematic representations of the experimental setup are shown in each panel. The retina from each mouse was used for staining whereas the other retina was used for gene expression analysis. Scale bars: 100 µm. (**A**) P5 retinas labeled with IB4 (greyscale) and stained for ERG (green), including quantification of ERG+ endothelial cells per retina 24 hours, and quantification of *Erg* mRNA levels relative to *Actb*. (**B**) P5 retinas labeled with IB4 (greyscale) and stained for Ki67 and pHH3, including quantification of Ki67+ and pHH3+ cells detected in IB4+ vasculature 24 hours; Ki67+ pHH3-cells (red), Ki67+ pHH3+ cells (pink), and quantification of mRNA levels for *Mki67* (encoding Ki67) relative to *Actb.* (**C**) P5 retinas labeled with IB4 (greyscale) and stained for p21 (magenta), including quantification of p21+ cells per retina and quantification of mRNA levels of *Cdkna1* (encoding p21) relative to *Actb*. Data are shown as mean ± SD. Each data point represents one retina from one mouse (n = 4-8 per group). Gene expression results are reported as fold change relative to the average of the CreER-group. \**P* < 0.05; \*\**P* < 0.01; ns: not significant (*P* > 0.05). Mann-Whitney U test.

To determine whether the reduction in ERG+ cells resulted from decreased proliferation, we examined P5 retinas after tamoxifen treatment on P2 and P4 by staining for IB4 together with two proliferation markers. Thus, we used an antibody for the proliferation marker Ki67, which is expressed during all active cell cycle phases [35], and antibodies specific for phospho-histone H3 (pHH3), which is detected in cells during the G2/M phase (but not in telophase; [36]). Quantitative analysis of the stained retinas (**Fig. 3B**) revealed a 49% reduction in Ki67⁺ endothelial cells in CreER+ compared to CreER-retinas, indicating that fewer endothelial cells were actively cycling after CreER activation (**Fig. 3B).** Consistent with the reduced number of Ki67⁺ endothelial cells in retinas 24 hours after the second tamoxifen dose, *Mki67* transcript levels were reduced by 36% in CreER+ compared to CreER-retinas (**Fig. 3B**). Agreeing with the reduced number of Ki67+ endothelial cells, we also observed 45% fewer pHH3+ endothelial cells in CreER+ compared to CreER-retinas (**Fig. 3B**). Contrasting the findings for endothelial cells, the number of Ki67+ non-endothelial cells was similar whereas the number of pHH3+ non-endothelial cells was increased in CreER+ versus CreER-retinas (**Fig. S1D**). Together, these findings suggest that a subset of non-endothelial cells had arrested in mitosis after CreER activation, which was not seen for endothelial cells, which instead had stopped cycling without an obvious mitotic arrest.

As the above data suggest that CreER activation stalls endothelial cell cycle progression, we examined retinas for the expression of p21, which is a known regulator of both DNA replication and DNA damage repair [37] and was previously shown to be upregulated after *Cdh5*-CreER activation [38]. We therefore performed quantitative analysis of the whole-mount retinas stained for IB4 and an antibody specific for p21 retinas 24 hours after the second of two tamoxifen injections (**Fig. 3C**). We found that the number of p21⁺ endothelial cells was increased 4-fold in CreER+ compared to CreER-retinas (**Fig. 3C**). Strikingly, p21 upregulation was confined to the IB4+ area despite ubiquitous *Cagg*-CreER activation (**Fig. S1E**). Quantitative RT-PCR analysis corroborated that p21 was significantly upregulated in CreER+ compared to CreER-retinas after two doses of tamoxifen (**Fig. 3C**). No such transcriptional upregulation was observed in P5 retinas of vehicle-treated mice or mice that had received only a single tamoxifen dose on P4 (**Fig. 3C**). Nevertheless, we observed an increase in p21⁺ endothelial cells in CreER+ compared to CreER-retinas after a single tamoxifen dose (**Fig. S2**), which suggests that p21 upregulation initially occurred independently of transcriptional induction. A post-transcriptional mechanism is consistent with the known regulation of p21 protein stability through ubiquitin-mediated degradation pathways [37]. Together, these results imply that p21 protein levels rapidly increase in endothelial cells following CreER activation and precede overt vascular defects, such as fewer ERG+ cells or reduced vascular extension and branching. This early response identifies p21 as a sensitive marker of CreER-mediated endothelial toxicity.

### CreER toxicity and p21 upregulation occurs in angiogenic not quiescent endothelium

To determine whether CreER-induced cell cycle effects were also observed with endothelial-selective CreER activation, we next examined mice carrying the *Cdh5*-CreER transgene, for which endothelial CreER toxicity was first described [13]. Similar to *Cagg*-CreER mice, P5 CreER+ mice, which had received two tamoxifen injections on P2 and P4, had 22% fewer ERG+ cells, a 59% decrease in the number of pHH3+ endothelial cells, and a threefold increase in the number of p21+ endothelial cells compared to Cre-negative littermates (**Fig. 4A,B**). In contrast, the retinas of 5/5 adult CreER+ mice treated with three consecutive tamoxifen injections (1 mg per injection every 24 hours) exhibited no obvious changes in vascular density compared to CreER-littermates, and p21+ endothelial cells were not observed (**Fig. 4C**) Similar findings were observed when CreER was activated in *Cagg*-CreER adult mice (data not shown). Together, these findings suggest that both ubiquitous and endothelial-specific CreER activation impairs endothelial cell proliferation following p21-induced cell cycle arrest, likely as a response to off-target Cre endonuclease activity.

**Fig. 4.**
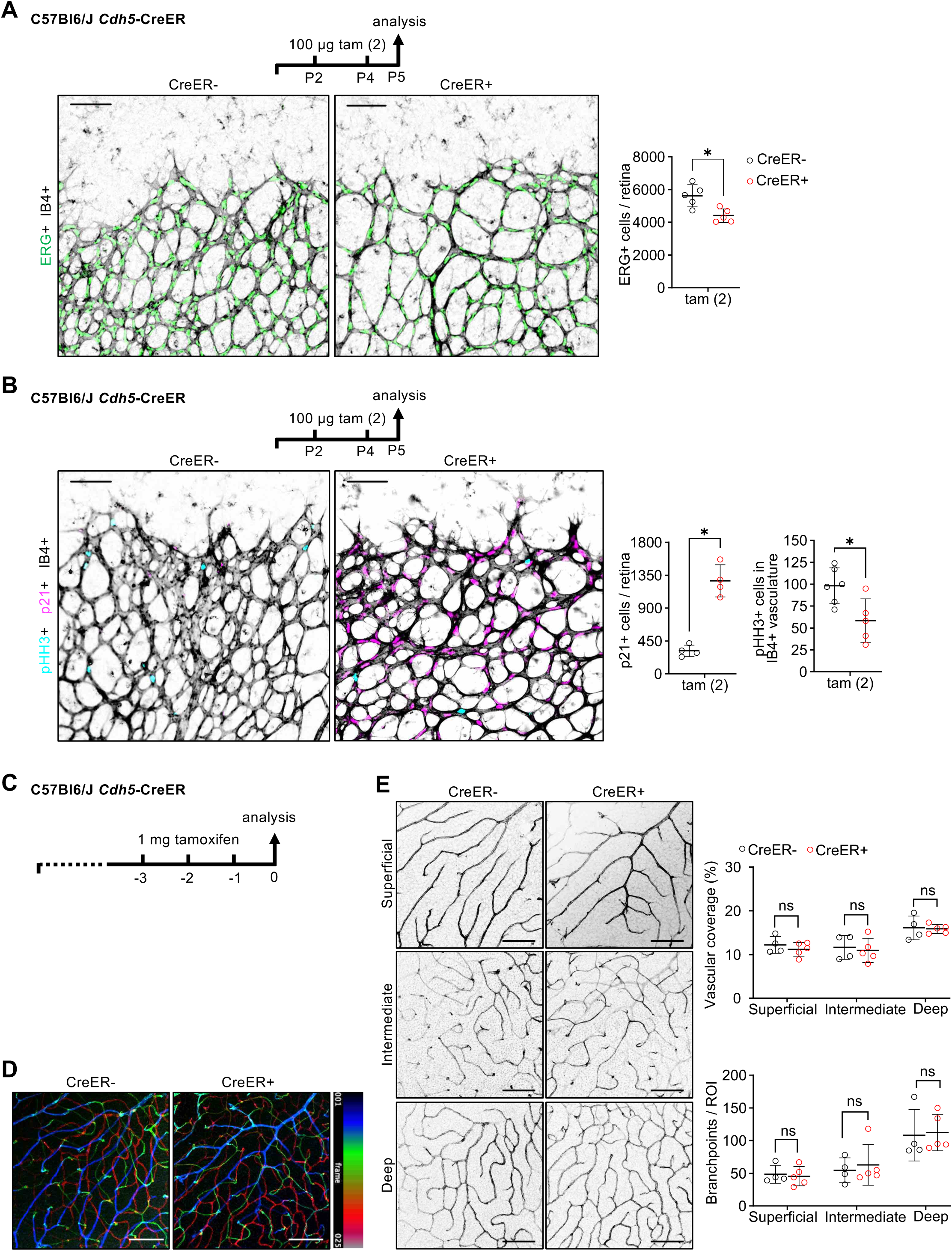
Reduced vascular growth and p21 upregulation in angiogenic but not in quiescent endothelium after *Cdh5*-CreER activation. (**A, B**) *Cdh5*-CreER+ and CreER-littermate pups on a C57BL6/J background were treated with 100 µg tamoxifen on P2 and P4, referred to as the tam (2) condition, and their retinas were analyzed on P5. Schematic representations of the experimental setup are shown in each panel. (**A**) P5 retinas labeled with IB4 (greyscale) and stained for ERG (green), including quantification of ERG+ endothelial cells per retina. (**B**) P5 retinas labeled with IB4 (greyscale) and stained for p21 (magenta) and pHH3 (cyan), quantification of p21+ cells per retina and pHH3+ cells detected in IB4+ vasculature per retina. Scale bars: 100 µm. (C) Schematic representation of the experimental setup of adult *Cdh5*-CreER+ and CreER-mice on a C57BL6/J background, which were treated with 1 mg tamoxifen daily for three consecutive days and analyzed 24 hours after the last injection. **(D)** Representative projection of confocal z-stack images from whole-mount retinas of CreER+ and CreER-mice stained with IB4; with vascular plexi resolved by pseudo-coloring according to retinal depth (25 z-slices; blue: superficial, green: intermediate, red: deep). **(E)** Representative images of the three IB4+ retinal plexi, shown in greyscale, extracted from the confocal z-projection shown in (**D**) including quantification of vascular coverage and branchpoints per region of interest (ROI) between an artery and vein in each plexus (n = 4-5 per group). Scale bars: 100 µm. Data are shown as mean ± SD. Each data point represents one ROI for one retina from one mouse. \**P* < 0.05; ns: not significant (*P* > 0.05). Mann-Whitney U test.

## Discussion

CreER-loxP mutagenesis in the mouse has considerably advanced our understanding of vascular biology and remains the primary approach for investigating angiogenesis regulators. For example, the spatiotemporal control of gene recombination in the developing retina has enabled pivotal insights into key regulatory pathways such as VEGF, TGFB and notch signaling, as well as their intricate cross talk during angiogenesis (e.g., [8,7,5,39,11]). However, recent studies have highlighted a significant caveat with this widely used system. Specifically, we and others have shown that endothelial cells in the angiogenic retina are highly susceptible to the toxic effects of CreER activation independently of targeted gene deletion [13,38], thereby creating risk that the interpretation of angiogenesis effects in knockdown studies may be inaccurate unless CreER toxicity is accounted for. Here, we demonstrate that CreER activation does not affect the quiescent endothelium and that Cre-induced vascular defects in the angiogenic retina are attributable to reduced endothelial proliferation. Importantly, our findings also reveal that CreER toxicity is not mitigated by the presence of loxP sites and that the angiogenic endothelium exhibits a capacity for recovery over time.

Ubiquitously expressed transgenes have been used to investigate the angiogenic effects of molecules expressed in endothelial cells (e.g., [16,9,40,14,15]). These transgenes are also valuable for reporting on angiogenic requirements of molecules secreted by other cell types in the vascular environment that affect endothelial cell behavior. While expected, our study shows that CreER activation has toxic effects on endothelial cells also when using a ubiquitous promoter, thereby reinforcing the importance of considering CreER-associated toxicity as a confounding variable in angiogenesis studies. Furthermore, we have made two additional notable observations, which suggest that endothelial-selective CreER activation should be prioritized when investigating potential angiogenesis regulators in endothelial cells, in order to minimize systemic confounding factors that arise from using a ubiquitously expressed CreER transgene. First, we detected reduced postnatal weight gain following *Cagg*-CreER activation across all tamoxifen doses tested, which was not observed for endothelial-specific CreER activation [13]. This effect was also observed when pups were assessed two weeks after tamoxifen discontinuation, when vascular density had normalized. Widespread CreER activation therefore imposes a broader burden on early postnatal development that cannot be attributed to endothelial toxicity alone, and suggests that ubiquitously expressed CreER should be avoided for CreER expression when studying endothelial genes. Second, activating a ubiquitously and highly expressed CreER transgene impaired vascular extension even at the lowest tamoxifen dose tested (50 µg), which falls into the lower range commonly reported in retinal angiogenesis studies. Nevertheless, the use of ubiquitous promoters to drive CreER expression remains necessary for targeting candidate pro-angiogenic modulators expressed by non-endothelial cells and will therefore require appropriate controls to account for endothelial toxicity [4].

We also found that the repeated delivery of even low doses of tamoxifen to activate CreER becomes increasingly toxic and therefore does not provide a reliable strategy to mitigate CreER-associated toxicity. Under our experimental conditions, a single tamoxifen injection had no detectable impact on retinal endothelial cell number 24 hours later; however, two injections caused a ∼35% reduction in endothelial cell density 24 hours after the second injection. Notably, the most commonly employed protocol for gene targeting in retinal angiogenesis studies involves three or four consecutive tamoxifen injections [3]. Our findings suggest that such repeated injections may exacerbate CreER-associated toxicity beyond the effect we have observed with two injections, although the magnitude of the effect will likely depend on CreER expression levels. In this context, a better understanding of the molecular mechanisms that underlie CreER toxicity for endothelial cells may help to identify dosing thresholds that have fewer off-target effects without compromising recombination of floxed target genes.

To determine which angiogenesis-relevant endothelial cell functions are affected by CreER activation, we considered previous reports showing that CreER activation lowers proliferation rates in mouse embryo fibroblasts and mouse keratinocytes *in vitro* [21]. Of note, we established that vascular defects can be detected earlier than previously reported [13] and shown here (**Fig. 1**); specifically, we observed a lower number of ERG+ cells already on P5, two days before obvious angiogenesis defects were detected on P7. Using Ki67 and pHH3 staining, we attributed this early growth defect to reduced endothelial cell proliferation after CreER activation. By contrast, CreER activation in the quiescent adult vasculature did not affect vascular density, thereby suggesting that endothelial cell proliferation is an angiogenic parameter susceptible to CreER toxicity. Consistent with our findings in retinal endothelium, a prior study using a Cx3Cr1-CreER transgene demonstrated that CreER activation adversely affected microglia proliferation in early postnatal brains, whereas no change in the microglia number was detected when Cx3Cr1-Cre was activated in adult mice [41]. Similarly, activating the ubiquitous *Rosa26*-CreER transgene reduced the proliferation of immature hematopoietic cells in the thymus and spleen of adult mice [42], further reinforcing the notion that rapidly proliferating cells are more vulnerable to the toxic effects of CreER activation.

According to our results, and those reported in a prior study [38], p21 upregulation is an early response to CreER activation during retinal vascularisation. Although p21 is a well-known effector of the DNA damage response and also serves as a cell cycle regulator [37], our experiments do not directly demonstrate causality between DNA damage and reduced endothelial proliferation. Nevertheless, the observed p21 upregulation is consistent with cell cycle arrest as a response to DNA damage. Considering that pseudo-loxP sites are unpaired, their cleavage is expected to cause single stranded breaks [17]. In non-proliferative cells, such breaks can be readily repaired with the opposing DNA strand as a template. By contrast, the single-stranded DNA present in S-phase is expected to be more vulnerable to DNA damage than double-stranded DNA and would require non-homologous end joining for repair. Supporting a model in which cells in S-phase are particularly vulnerable to CreER toxicity due to DNA damage, CreER activation did not upregulate p21 in endothelial cells of the quiescent retinal vasculature. Therefore, it is conceivable that CreER activation-induced p21 upregulation could stall cell cycle progression to allow for single stranded DNA repair. Restoring endothelial health by imposing a break in the cell cycle, including in DNA synthesis, might therefore allow p21 to pave the way for the vascular normalization we observed two weeks after tamoxifen discontinuation. Nevertheless, we cannot exclude that some endothelial cells may also die after p21 upregulation, because it is known to trigger apoptosis in other cell types in response to unrepaired DNA damage [37].

Notably, p21 upregulation was restricted to endothelial cells, not only after endothelial-selective CreER activation but also after ubiquitous CreER activation in the retina. Consistent with a protective role for p21 in promoting vascular resilience to CreER-induced stress, analysis of pHH3 and Ki67 staining in the retina revealed fewer cycling endothelial cells after CreER activation, without evidence of mitotic arrest. In contrast, a large number of non-endothelial cells, which showed no p21 upregulation, appeared to have stalled in mitosis, potentially indicating a cell cycle block. Importantly, p21 upregulation in retinal endothelial cells was detectable as early as 24 hours after a single tamoxifen dose and preceded the detection of effects on cell cycle progression. These findings raise the question why endothelial cells in the retina are uniquely primed to mount a rapid p21-mediated response to CreER activation. A plausible explanation might arise from the prior finding that p21 serves a physiological function in retinal vascular development. Specifically, it has been reported that p21 is selectively upregulated in a subset of endothelial cells at the vascular front to balance endothelial cell sprouting and proliferation in the context of high mitogenic stimulation [43]. It is therefore conceivable that CreER-mediated p21 upregulation and its associated function in regulating endothelial cell proliferation co-opts a physiological mechanism that normally operates at the angiogenic front at low levels, and thus serendipitously confers vascular resilience.

In summary, our results support the concept that CreER toxicity is a significant confounder in studies of postnatal angiogenesis and reveal that these effects also occur when a ubiquitously expressed transgene is employed to target endothelial cells. These findings emphasize the need for tamoxifen-activated CreER controls to disentangle CreER-induced effects from gene-specific phenotypes due to loxP-mediated recombination. Although our evidence suggests that quiescent vasculature in adult animals appears less affected by CreER toxicity, further work is required to understand implications for mouse models of adult neovascular diseases, in which endothelial cell proliferation is reactivated, for example, in choroidal neovascular disease, tumor angiogenesis, or ischemic vascular diseases such as peripheral artery disease or myocardial infarction.

## Non-standard Abbreviations and Acronyms

(EC): Endothelial Cell
(CreER): Cre recombinase estrogen fusion protein with ligand binding mutation

## Acknowledgements

We thank the staff of the Biological Resources and the Microscopy Facility at the UCL Institute of Ophthalmology for their valuable assistance; James Brash and Laura Denti for technical help in the initial phase of this study; and Tiago V. Pereira for statistical analysis assistance.

## Methods

### Animals

All procedures were conducted in accordance with the Institutional Animal Welfare Ethical Review Body and the United Kingdom Home Office guidelines under the Animals (Scientific Procedures) Act 1986. All mice were bred at the UCL Institute of Ophthalmology’s Biological Resources Unit and housed in individually ventilated cages at 20-24°C ambient temperature, 55±10% relative humidity, and a 12-hour light/dark cycle. All mice had access to a standard diet and water ad libitum. In this study, Tg(CAG-cre/Esr1*)5Amc (*Cagg*-CreER) (MGI:212767) [31] or Tg(Cdh5-cre/ERT2)#Ykub (*Cdh5*-CreER) (MGI:5705396) [11] males a C57BL6/J background were crossed with wild-type females a C57BL6/J background to generate mice carrying or lacking ubiquitous and endothelial cell-specific tamoxifen-inducible CreER, respectively. In some experiments, *Cagg*-CreER males were crossed with GT(*Rosa)26Sor^tm14(CAG-tdTomato)Hze^* (*Rosa26^tdTom^*) (MGI:3813512) females on a mixed genetic background (C57BL6/J:129/Sv).

### Tamoxifen treatments

Tamoxifen powder (Sigma, T5648) was dissolved in vegetable oil (peanut oil, Sigma P2144, or corn oil, Sigma C8267) by agitation at 37°C for 2 hours. To activate CreER during the perinatal period, mice received two intraperitoneal injections of 25 µL tamoxifen (2, 4 or 6 mg/mL) on postnatal day (P) 2 and P4, and eyes were collected on P5, P7 or P21. For experiments assessing the effects of a single tamoxifen injection, mice received one 25 µL tamoxifen (4 mg/mL) intraperitoneal injections on P4, and eyes were collected on P5. To activate CreER in adult mice (6-8 weeks old), 100 µL tamoxifen (10 mg/mL) was administered via intraperitoneal injections once daily for three consecutive days, and eyes were collected 24 hours after the final injection. Littermate animals lacking CreER received intraperitoneal tamoxifen or vehicle (corn oil) injections to serve as controls. Both male and female mice were included in all analyses.

### Whole-mount retina staining

Eyes were enucleated at the indicated time points and fixed in 4% formaldehyde in PBS for 10 minutes at room temperature (RT). Retinas were immediately micro-dissected and prepared for flat mounting by making four radial incisions. Freshly dissected retinas were stored in 96-well plates at -20°C in 100% methanol until staining. Before immunostaining, retinas were rehydrated in PBS and permeabilized in PBS containing 0.5% Triton X-100 (PBST) for 30 minutes at RT. P5 retinas were incubated in blocking solution comprised of 10% heat-inactivated normal goat serum and 1% bovine serum albumin (Sigma-Aldrich) in PBST for 1 hour at RT. Primary antibodies were diluted 1:400 in blocking solution and incubated for 90 minutes at RT; the following primary antibodies were used: anti-phospho-histone H3 (anti-phH3; H9908, Sigma-Aldrich), anti-Ki-67 (ab16667, Abcam), anti-ERG (ab92513, Abcam) and anti-p21 (ab188224, Abcam). Following four 15-minute PBS washes, retinas were incubated with fluorescein-labeled G. simplicifolia lectin (GSL I–BSL I; FL-1101-5, Vector Labs) and the following secondary antibodies, diluted 1:200 in PBS for 1 hour at RT: Alexa Fluor® 647 AffiniPure® Fab Fragment Donkey Anti-Rabbit IgG (H+L) (711-607-003, Stratech Jackson), Cy™3 AffiniPure® F(ab’)₂ Fragment Donkey Anti-Rat IgG (H+L) (712-166-150, Stratech Jackson), Cy™3 AffiniPure® F(ab’)₂ Fragment Donkey Anti-Rabbit IgG (H+L) (711-166-152, Stratech Jackson). After four 15-minute washes in PBS, retinas were mounted on glass slides using VECTASHIELD^®^ Vibrance™ Antifade Mounting Medium (H-1700-10, 2BScientific). P7 and P21 retinas were blocked in 10% serum-free protein block (DAKO; X0909, Agilent) in PBST for 1 hour at RT. Biotinylated IB4 (L2140, Sigma) was diluted 1:200 in the blocking solution and incubated overnight at 4°C with gentle rocking. After washing three times for 10 minutes in PBS, retinas were incubated for 2 hours at RT with streptavidin conjugated to Alexa Fluor® 488, 647 or 350 (S11223, S21374 or S11249, respectively, Thermo Fisher Scientific), diluted 1:200 in the blocking solution. Stained retinas were washed three times for 10 minutes in PBS and mounted on glass slides using Fluoromount-G™ Mounting Medium (00-4958-02, Thermo Fisher Scientific).

### Image acquisition and analysis

IB4-stained P7 retinas were imaged using an SZX16 fluorescent stereomicroscope (Olympus) equipped a C4742-95 camera (Hamamatsu). Retinal radius and retinal vascular extension were measured in each leaflet using the FIJI (NIH Bethesda) and averaged by retina. Retinal radius was defined as the distance from the optic disc to the retinal margin, whereas retinal vascular extension was defined as the distance from the optic disc to the front of the vascular network divided by the retinal radius. Vascular branchpoints were quantified in 4 fields of view per retina (150 x 150 pixel each) behind the angiogenic front in a region between an artery and a vein using Angiotool v.0.9226 (NIH Bethesda) and averaged by retina. IB4-stained P5 retinas co-stained for ERG, pHH3, Ki-67 and P21 imaged using a widefield inverted Ti2 fluorescent microscope (Nikon) equipped with a DS-Qi2 camera (Nikon). The entire leaflet (or whole retina for ERG) was analyzed using FIJI. Background fluorescence was subtracted using a rolling ball algorithm with a radius of 10 pixels for ERG and p21, 30 pixels for Ki-67, and 50 pixels for pHH3 and IB4. The number of positively stained cells was automatically quantified using the “Analyze Particle” function in FIJI after fluorescent signal thresholding and then confirmed by visual inspection of each image. Co-localization of pHH3+ or Ki-67+ cells with IB4 was assessed manually. The values for each leaflet were summed to obtain the value for that retina. Images of IB4-stained P21 retinas were acquired with a Stellaris 5, confocal microscope equipped with LAS X software (Leica) and comprised a 581 µm^2^ region between an artery and a vein, with the scan beginning at the surface of the retina and extending through the ganglion cell layer into the outer plexiform layer. Z-projections of the vasculature were generated using a temporal color-coding method in FIJI. Confocal slices containing the superficial, intermediate or deep vascular plexus were extracted from each z projection and processed separately for quantification of vascular coverage; the number of branchpoints relative to the retinal area was determined using Angiotool v0.9226 (NIH Bethesda) in four regions of interest (ROI) for P21 and one ROI for adults. When four ROIs were acquired per retina, it was one for each leaflet, and the values were averaged for each plexus in each retina.

### Gene expression analysis

Total RNA was isolated from P5 retinas using the RNeasy Mini Kit (Qiagen Inc.) and reverse-transcribed using the Superscript IV reverse transcription kit (Thermo Fisher Scientific). cDNA was analyzed by real-time PCR on the QuantStudio 6 Flex Real-Time PCR system (Applied Biosystems) using the SYBR Green master mix (Applied Biosystems) and oligonucleotide primer pairs specific for *Erg* (forward 5’-CCGGATACTGTGGGGATGAG-3’ and reverse 5’-TCTGCGCTCATTTGTGGTCA-3’), *Cdkn1a* (forward 5’-TCGCTGTCTTGCACTCTGGTGT-3’ and reverse 5’-TCGCTGTCTTGCACTCTGGTGT-3’), *mKi67* (5’-GAGGAGAAACGCCAACCAAGAG-3’) and *Actb* (forward, 5’-CACCACACCTTCTACAATGAG-3’ and reverse 5’-GTCTCAAACATGATCTGGGTC-3’). Expression of each target gene was normalized to *Actb* levels, and quantification was performed using the 2^-ΔΔCT^ method [44]. Values are expressed as fold change in the CreER+ relative to the CreER-control group.

### Statistical analysis and reproducibility

One retina from one mouse was considered one biological replicate, whereby both retinas were used for different analyses (e.g., one for staining and the other for gene expression). We analyzed 4 different fields of view from each retina and averaged the values to obtain the value for that sample. Blinding was used during treatment and outcome assessment, with genotypes being disclosed only during statistical analysis. Samples or data points were excluded only in the case of technical equipment or human error that caused a sample to be poorly controlled. Statistical analyses were performed using Prism 10 (GraphPad Software Inc.) or Stata (version 18, StataCorp, College Station, Texas, USA). Data are shown as means ± SD. A non-parametric Mann-Whitney test was performed for all analyses. Interaction between tamoxifen doses and sex was tested by comparing the goodness-of-fit of two linear regression models: a reduced model including only the main effects of treatment and sex, and a full model that also included the treatment-by-sex interaction term. We then used a likelihood-ratio test to test if the full model offered a better fit to the data than the reduced model. Significance was established at *P*< 0.05. *P* values are indicated in each Figure as \**P*< 0.05, \*\**P*< 0.01, \*\*\**P*< 0.001, \*\*\*\**P*< 0.0001. At least two independent litters were used for each analysis to ensure reproducibility and robustness of findings. Sample size estimates for vascular extension as the primary outcome were based on the number of animals needed to observe an effect size of 2.0 standard deviation units (a mean difference of 0.1 and a common SD of 0.05, based on prior published data [13]; using calculations with a 2-sided alpha 0.05 and power 80%, the required sample size was calculated to be seven mice per group.

