## Supplemental figures for "CreER activation transiently impairs angiogenesis by slowing endothelial proliferation"

#### Supplemental data

##### Supplemental Fig. S1

**Increased number of non-endothelial cells in mitosis and endothelial cell selectivity for p21 upregulation after *Cagg*-CreER activation**

(**A**) Schematic representation of the tamoxifen treatment of *Cagg*-CreER+ and CreER- pups carrying or not carrying the *Rosa26*$^{tdTom}$ reporter, analyzed on P7 following tamoxifen administration of the indicated doses on P2 and P4 (n = 5-14 pups per group). In the graphs, red and blue circles represent the values from tdTom+ and tdTom- retinas, respectively.

(**B**) Schematic representation of the experimental setup and body weight of *Cagg*-CreER+ and CreER- littermate pups on a C57BL6/J background, treated with 100 µg tamoxifen on P4, referred to as the tam(1) condition, and then analyzed on P5 (n = 9-12 per group).

(**C-E**) *Cagg*-CreER+ and CreER- littermate pups on a C57BL6/J background were treated with 100 µg tamoxifen on P2 and P4, referred to as the tam (2) condition, and then analyzed on P5. Schematic representation of the experimental setup is shown in each panel. (**B**) Body weight (n= 16 per group).

(**C**) Quantification of Ki67+ and pHH3+ cells detected outside of the IB4+ vasculature per retina 24 hours after the second tamoxifen dose using images generated for endothelial cell analysis in Fig. 3

(**D**). Representative images of P5 retinas stained with IB4 (blue) and antibodies for pHH3 (green) and p21 (red) to illustrate endothelial selectivity of p21 upregulation. Scale bars: 100 µm.

Data are shown as mean ± SD. Each data point represents one mouse. ***P* < 0.01; ****P* < 0.001; *****P* < 0.001; ns: not significant (*P*>0.05). Mann-Whitney U test.

##### Supplemental Fig. S2

**No p21 upregulation in angiogenic endothelium due to vehicle treatment but 24 hours after a single tamoxifen injection**

(**A, B**) *Cagg*-CreER+ and CreER- littermate pups on a C57BL6/J background were treated with vehicle on P2 and P4 (**A**) or with 100 µg tamoxifen on P4, referred to as the tam (1) condition, and then analyzed on P5. Retinas were labeled with IB4 (greyscale) and stained for p21 (magenta). Scale bars 100 µm.

### Supplemental Figure S1

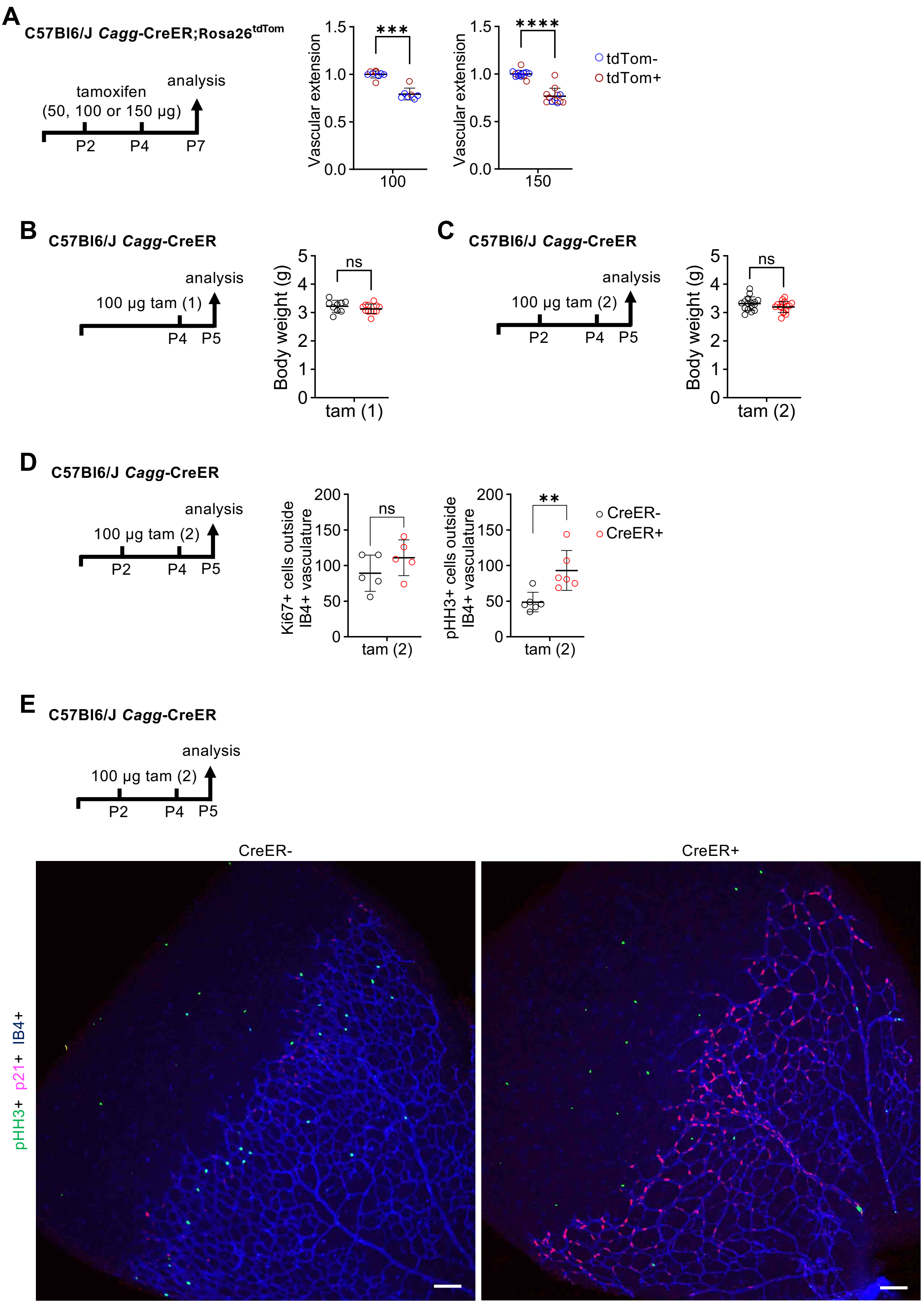

### Supplemental Figure S2

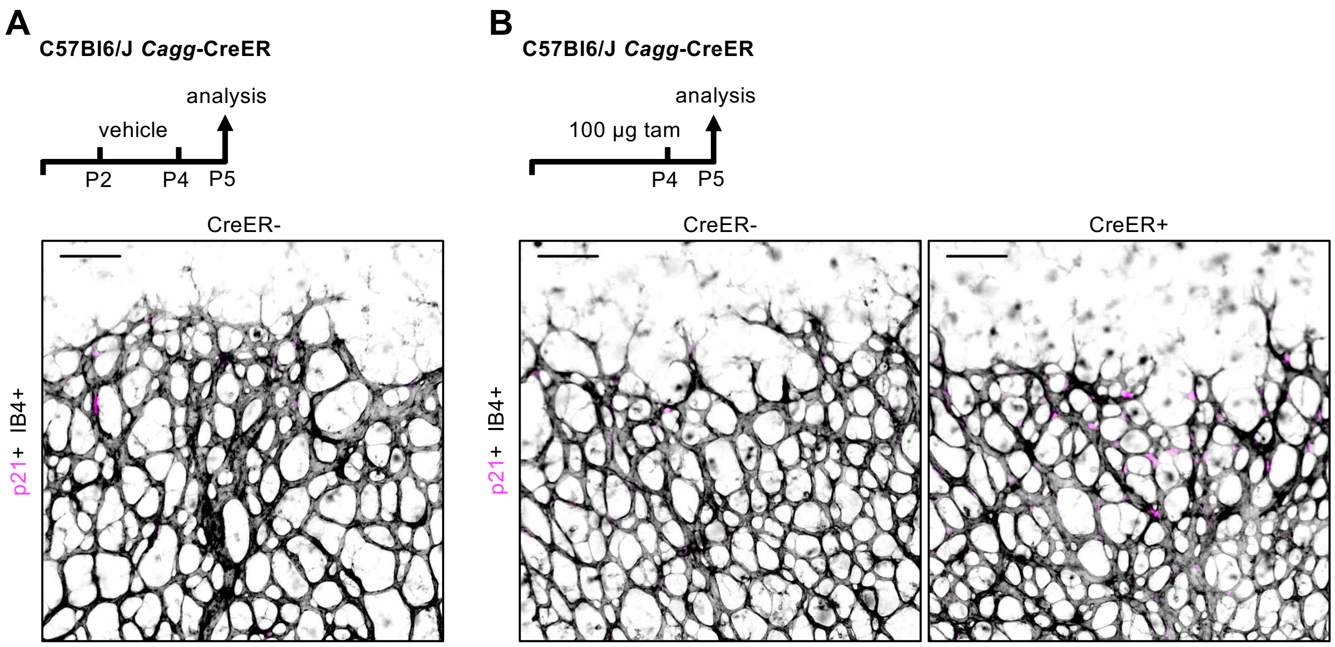
